# Label-free quantification in the Crux toolkit

**DOI:** 10.1101/2025.04.21.649897

**Authors:** Frank Lawrence Nii Adoquaye Acquaye, Bo Wen, Charles E. Grant, William Stafford Noble, Attila Kertesz-Farkas

## Abstract

Ultimately, most tandem mass spectrometry (MS/MS) proteomics experiments aim to not just detect but also quantify the proteins in a given complex sample. Here, we describe an extension to the Crux MS/MS analysis toolkit to enable label-free quantification of peptides. We demonstrate that Crux’s new quantification command, which is modeled after the algorithms implemented in the widely used FlashLFQ software, is both efficient and accurate. In particular, we achieve a 1.9-fold speedup while reducing the memory usage by 26%. The new crux-lfq command is available in Crux v4.3.

## 1 Introduction

Three general classes of methods have been developed to extract quantitative measurements from proteomics tandem mass spectrometry (MS/MS) data acquired using data-dependent acquisition. Perhaps the simplest approach involves counting the number of times that peptides associated with a given protein are detected in a particular MS/MS run. These spectral counting methods yield only “semi-quantitative” estimates of protein abundance. A second, increasingly popular category of methods involve labeling peptide molecules with various types of tags—iTRAQ or TMT—and using the fragment ion intensities associated with these tags as a quantitative measure of abundance. These tag-based approaches yield much more accurate quantitative measurements; however, the quantitative values often suffer from ratio compression [1], and the tagging procedure adds complexity and expense to the assay. The third category, intermediate in complexity between these two approaches, consists of label-free quantification (LFQ) methods. These methods involve detecting the presence of a particular peptide species based on a fragmentation (MS2) spectrum and then computing the abundance based on peak intensities in the associated precursor (MS1) spectra.

Many LFQ software tools are available and in widespread use (Table 1). Unfortunately, the most widely used tool, MaxQuant [2], is closed source. Because the Crux software [3] aims to provide a general toolbox for proteomics mass spectrometry analysis, we aimed to add an LFQ tool to Crux. Rather than inventing a new approach, we adopted the successful algorithm in FlashLFQ [4]. This algorithm has the advantage of being much faster than some competing methods, since it uses a hash table to match MS1 peaks on the fly to peptides detected in the MS2 data. Relative to FlashLFQ, the Crux implementation offers better interoperability with the other tools in Crux as well as a nearly two-fold speed-up relative to FlashLFQ.

**Table 1:** LFQ software tools. Citation counts are from Google Scholar as of March 31, 2025.

| Method | Year | Availability | Language | Citations | Citations since 2022 |
| --- | --- | --- | --- | --- | --- |
| MaxQuant [2] | 2008 | Closed source | C# | 5262 | 2109 |
| moFF [5] | 2016 | Open source | Python | 65 | 11 |
| FlashLFQ [4] | 2018 | Open source | C# | 104 | 58 |
| IonQuant [6] | 2020 | Closed source | Java | 305 | 290 |
| IceR [7] | 2021 | Source Available | R | 50 | 47 |
| Sage [8] | 2023 | Open source | Rust | 47 | 47 |
| CruxLFQ | 2025 | Open source | C++ |  |  |

## 2 Methods

### 2.1 Quantification algorithm

The CruxLFQ software implements the algorithmic approach pioneered by FlashLFQ [4]. The algorithm proceeds in three steps. First, the program reads a file containing peptide-spectrum matches (PSMs) and, for each distinct peptide sequence, calculates a corresponding isotope distribution. Second, the program reads the corresponding MS1 data into memory, storing each observed MS1 peak only if it matches the *m/z* of one of the PSMs from the first step. Finally, in the third step, the matches from the second step are refined by analyzing each one in the retention time dimension, looking for gaps, enforcing a tighter *m/z* threshold, and ensuring that the chromatogram exhibits a “peak-like” behavior.

### 2.2 Software implementation

The LFQ algorithm of CruxLFQ is implemented as the lfq command in the Crux toolkit [3, 9]. The software takes as input a list of spectrum files in mzML format and a PSM file in either of two tab-delimited formats, as produced by Percolator (http://percolator.ms) or Crux’s assign-confidence command. CruxLFQ is implemented in C++, is open source with an Apache license, and is distributed with pre-compiled binaries for Windows, MacOS and Linux (http://crux.ms). User support is provided via a Github issue tracker (https://github.com/crux-toolkit/crux-toolkit/issues).

### 2.3 Benchmarking dataset

For benchmarking, we use the IonStar collection of MS/MS runs from human Panc-1 cells mixed with DH5*α E. coli* (ProteomeXchange identifier PXD005590) [10]. This dataset was also used as a benchmark in the FlashLFQ paper [4]. The dataset consists of 20 runs, corresponding to four replicates each of five different amounts (A:2, B:3, C:4, D:5, or E:6% by weight, representing 1.-, 1.5-, 2.-, 2.5-, or 3.-fold addition) of *E. coli* peptides added to a constant amount of human peptides.

### 2.4 Peptide detection and quantification

The analysis of raw MS/MS files involved the conversion to centroided mzML files and a subsequent search against a protein database containing both *Homo sapiens* and *Escherichia coli* proteins (Human ecoli trypsin 1501v uniprot sprot.fasta from PXD003881) using several tools. Tide version 4.3 was used for both CruxLFQ and FlashLFQ (version 1.0.559) analyses, with fixed carbamidomethyl (C) modifications and variable modifications, including oxidation (M) and acetylation (Protein N-term), while applying the Trypsin/P enzyme with a maximum of two missed cleavages and a precursor window of 20 ppm. In CruxLFQ, peptides were accepted at a 1% peptide-level FDR threshold based on percolator (for Figure 1) or the assign-confidence command from Crux (for Figure 2 and Supplementary Table 1) and then quantified via CruxLFQ. FlashLFQ also processed the results from Crux, specifically employing mzlib version 1.0.559 for quantification. Sage (version 0.14.7) was employed similarly, with lfq quantification enabled, and the same modification parameters were followed. Quantification using IonQuant (version 1.10.27) was performed through FragPipe (version 22.0) with the default workflow in FragPipe, MSFragger (version 4.1) [11] was used for peptide detection, MS1 quantification was enabled but the match between runs option was disabled, again adhering to the same modification criteria. MaxQuant (version 2.6.3.0) was set up with identical modifications and enzyme rules but specifically had the “Match between runs” setting disabled, capturing the distinct nuances of each tool in their approaches to peptide quantification. All other parameters of MaxQuant were set to default.

**Figure 1:**
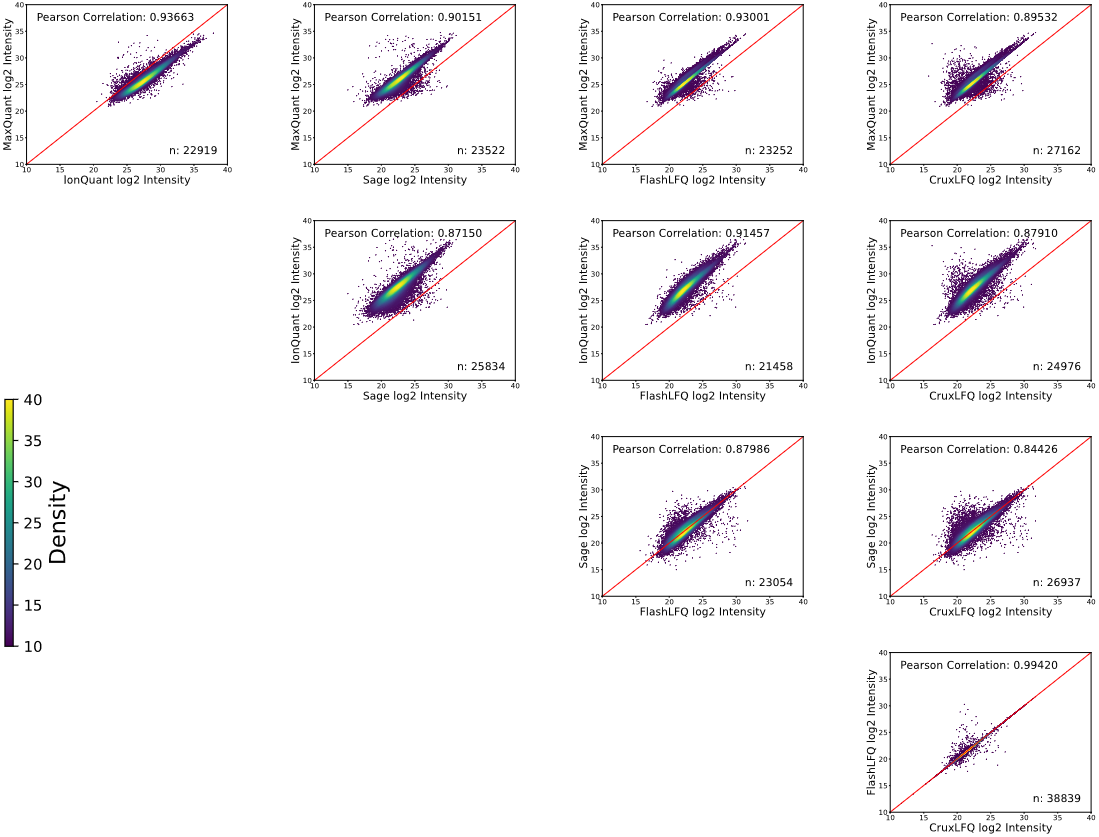
Comparison of five different LFQ tools. Each panel plots peptide log_2_ intensities as computed by two different LFQ tools, including MaxQuant, IonQuant, Sage, FlashLFQ and CruxLFQ. The data is from a randomly selected run from the human-*E. coli* dataset, specifically, the spectrum file B02_001_161103_ B1_HCD_OT_4ul.

**Figure 2:**
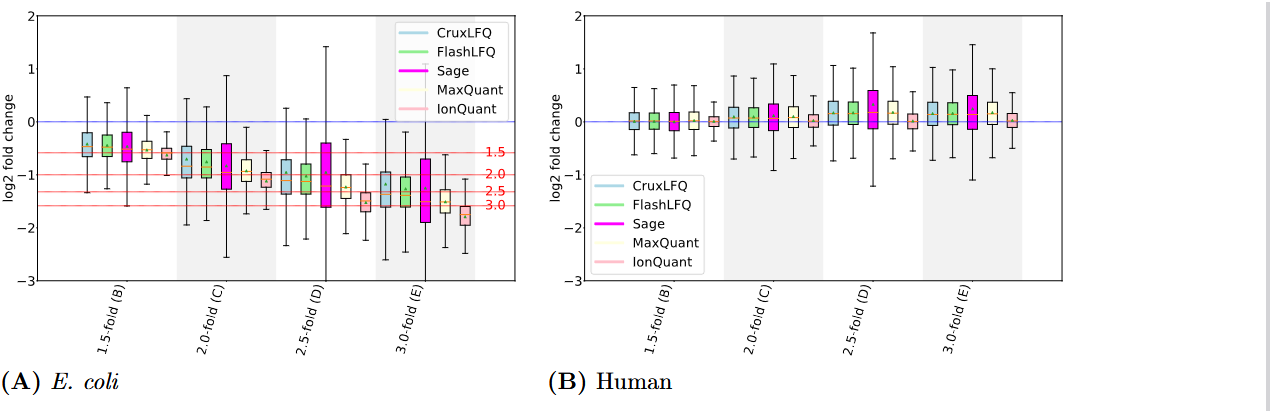
Trend comparison. **(A)** Distribution of log_2_ peptide fold-change values computed by various quantification methods on five varying amounts of *E. coli* proteins spiked into a human sample [10]. The *E. coli* protein amounts represent various fold-changes (A:1, B:1.5, C:2, D:2.5, E:3-fold changes). The bars show the log-fold change values of (A vs. B), (A vs. C), (A vs. D), and (A vs. E). Horizontal lines correspond to the expected log_2_ fold-change values. **(B)** Similar to panel **(A)**, except that instead of comparing various *E. coli* quantifications, we compare quantifications of human proteins between pairs of experiments. Because the amount of human protein was the same in each case, the expected log_2_ fold-change is zero.

The resulting quantification values were collected into one peptide-by-run matrix for each of the five analysis methods (Supplementary Table 1). All analyses except the MaxQuant and FragPipe search were carried out on a server equipped with an Intel Xeon CPU E5-2640 v4 2.40GHz processor with 20 cores, 128 GB DDR RAM, and storage capacity of 10 TB operated by Ubuntu 22.04.4 LTS OS.

## 3 Results

### 3.1 CruxLFQ results are consistent with existing LFQ tools

To validate CruxLFQ’s quantitative measurements, we began by comparing peptide-level intensities com- puted by FlashLFQ, MaxQuant, Sage, IonQuant and CruxLFQ on a common dataset. For this evaluation, we used a collection of 20 MS/MS runs derived from varying mixtures of human and *E. coli* proteins (four replicates and five different mixture ratios). Each run was searched using four different search engines: Tide, Sage, MSFragger and MaxQuant. When comparing two LFQ methods that rely on different search engines, only peptides that were confidently identified by both search engines and were assigned non-zero quantification values were considered. Scatter plots comparing the computed peptide intensities in one randomly selected run show strong correlation (Figure 1A), which is consistent with the average correlations computed across all 20 runs (Table 2). In particular, the correlation between CruxLFQ and FlashLFQ is very close to 1. The few differences arise because CruxLFQ calculates isotope distributions using code from the Hardlkor codebase [12].

**Table 2:** Pearson correlations between outputs of five different LFQ tools. Each entry is the average and standard deviation of the Pearson correlation between two LFQ tools, computed across 20 runs in the IonStar dataset. The underlying per-run correlations are provided in Supplementary Table 2.

|  | IonQuant | Sage | FlashLFQ | CruxLFQ |
| --- | --- | --- | --- | --- |
| MaxQuant | $0.93579 \pm 0.00151$ | $0.89838 \pm 0.00306$ | $0.92787 \pm 0.00578$ | $0.89053 \pm 0.01170$ |
| IonQuant | | $0.86934 \pm 0.00221$ | $0.91229 \pm 0.00231$ | $0.87796 \pm 0.00775$ |
| Sage | | | $0.87581 \pm 0.00229$ | $0.84022 \pm 0.00755$ |
| FlashLFQ | | | | $0.99554 \pm 0.00054$ |

### 3.2 Validation using multi-species spike-in data

For further validation, for each method we computed the *E. coli* and human protein quantification values in the IonStar dataset. In this data, five varying amounts of *E. coli* lysate (representing fold-changes of 1, 1.5, 2, 2.5 and 3) were spiked into a constant amount of human lysate [10]. The distribution of the log fold-change of quantification values (Figure 2) suggests that CruxLFQ produces log fold change values that are close to the ground truth fold changes for *E. coli* proteins (panel A) while maintaining a fold-change close to zero for pairs of human samples (panel B). The CruxLFQ quantification largely agrees with the results of the other methods, though IonQuant seems to consistently underestimate the *E. coli* fold change, whereas other methods consistently overestimate it. IonQuant also exhibits the lowest variance, relative to other methods. On the human peptides, IonQuant appears to perform the best overall.

### 3.3 CruxLFQ is efficient in both speed and memory usage

We next investigated the relative speed and memory requirements of CruxLFQ and FlashLFQ. To do so, we ran both tools on nested subsets of the 20 runs in the IonStar dataset (of sizes 1, 2, 4, 8, 16 and 20 runs). We repeated this procedure for three different random orderings of the twenty runs and the computed the mean runtime and RAM usage for each tool.

The results (Figure 3) show that CruxLFQ is consistently more efficient, in both time and memory usage, than FlashLFQ. For this dataset, CruxLFQ is 1.85 times faster than FlashLFQ on average. FlashLFQ takes approximately 4.37 minutes to analyze each raw file in the IonStar dataset, whereas CruxLFQ takes 2.36 minutes. Furthermore, for analyses of datasets consisting of 4–20 runs, FlashLFQ consistently requires, on average, 22.8% more RAM than CruxLFQ.

**Figure 3:**
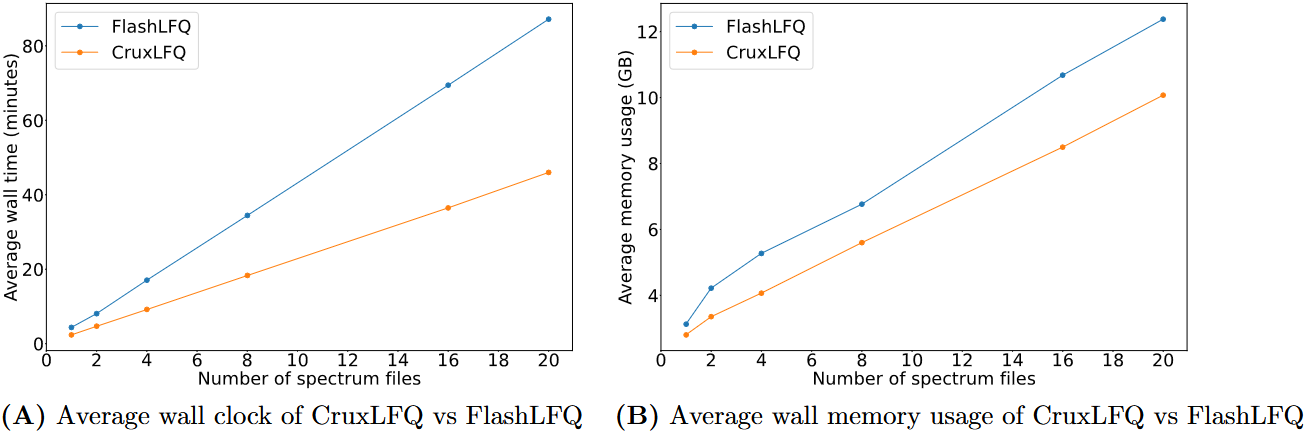
CruxLFQ vs FlashLFQ bench-marking. **(A)** Wall clock time of CruxLFQ and FlashLFQ as a function of the size of the dataset, averaged over three randomized sets of spectrum files. **(B)** Memory usage of CruxLFQ and FlashLFQ as a function of the size of the dataset, averaged over three randomized sets of spectrum files.

## 4 Discussion

Direct comparisons of quantification values produced by five different LFQ tools suggests that the values are largely consistent from one tool to the next. Furthermore, the corresponding analysis of fold-change values on the IonStar benchmark dataset suggests that, while all of the methods yield roughly accurate quantification values, no single method is clearly outperforming the others. In general, all four methods achieve a mean log-fold change that is within one standard deviation of the expected log fold-change, across a range of fold-change values. Better understanding why some methods seem to consistently over-predict the log fold-change while others occasionally over- or under-predict would be interesting but is challenging because several of these tools are closed source.

Adding label-free quantification functionality to Crux expands the utility of the toolkit, but there is still more work to be done. First, we have recently demonstrated how to carry out rigorous error rate control when propagating peptide identities between runs [13]. We therefore plan to implement this “PIP-ECHO” procedure as an option in CruxLFQ. We also aim to add a command to enable tandem mass tag and isotope-based quantification.

## Acknowledgment

This research was supported by National Science Foundation award 2245300 and in part through computational resources of HPC facilities at HSE University [14].

## Author contributions

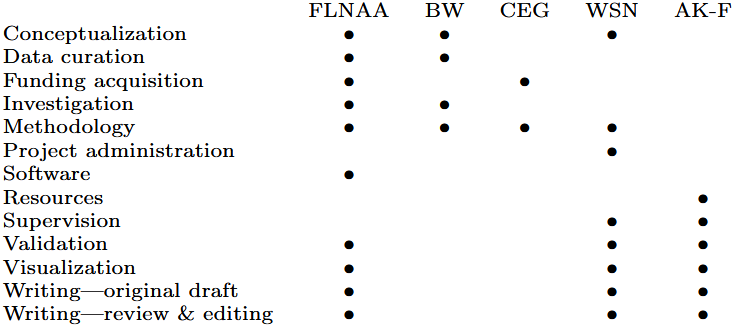

## Competing financial interests

The authors declare no competing financial interests.

## Availability

The new CruxLFQ is available in the Crux toolkit at https://crux.ms version 4.3 Apache licensed source code is available, and pre-compiled binaries are provided for Windows, MacOS and Linux.

## Notes

http://crux.ms

### Competing Interest Statement

The authors have declared no competing interest.

